# Succession of embryonic and intestinal bacterial communities of Atlantic salmon

**DOI:** 10.1101/128066

**Authors:** Jep Lokesh, Viswanath Kiron, Detmer Sipkema, Jorge M.O. Fernandes, Truls Moum

## Abstract

Host-associated microbiota undergoes continuous transition to achieve a stable community, and these modifications are immediately initiated from the birth of the host. In the present study, the succession of early life (eyed egg, embryo, and hatchling stages) and intestinal (the whole intestine at the early freshwater stages and the distal intestine at the late freshwater and seawater stages) bacterial communities of Atlantic salmon (*Salmo salar*; a prominent farmed fish) were studied using a 16S rRNA gene (V3 region) amplicon sequencing technique.

Stage-specific bacterial community compositions and the progressive transitions of the communities were evident in both the early life and the intestine. The embryonic communities were relatively less diverse, but after hatching the diversity increased significantly. A marked transition of the intestinal communities also occurred during the development. The most abundant functional pathways associated with the different stages were not affected by the transition of the community composition A perceptible transition in the community composition occurred during the development of Atlantic salmon. The transition generally did not alter the core functions of the community. Hatching and transfer to seawater are the key events that affect the bacterial diversity and community composition. The contribution of host-derived factors and environment in shaping the bacterial communities need to be confirmed through further studies.

## Introduction

All animals are born into a microbe-rich environment, and the host establishes a symbiotic relationship with its microbial community. Such symbiotic relationships play vital roles in the physiological functions of the host [1-4]. The unstable and compositionally variable microbiota associated with early life undergoes continuous transitions (from the first few days to the first few years of life) to achieve a compositional profile resembling that of adults [5-8]. Although the transition from the infant to the adult microbiome of humans is relatively well documented, the transition during the ontogeny of fish and the establishment of their microbial communities are relatively less explored. Studies on the microbiota of larval Atlantic cod (*Gadus morhua*) and killifish (*Kryptolebias marmoratus*) [9, 10], and the intestinal microbial communities during the ontogeny of zebrafish (*Danio rerio*) [11, 12] have shed light on the functional importance and transformation of the early life communities in fish. Furthermore, the transition of the bacterial composition during the ontogeny of wild Atlantic salmon belonging to different cohorts was described by Llewellyn *et al*. [13]. However, the general pattern of transition of the microbial communities in fish could be better explored if these fish belonged to a single cohort and were maintained under controlled conditions.

Atlantic salmon is an anadromous fish of high commercial value. In aquaculture production systems, embryos and larvae maintained in freshwater are immediately offered feeds after yolk sac absorption (~7-8 weeks post-hatching) and further reared in freshwater until these fish become smolts (a developmental stage that enables the fish to adapt to its physiological needs in seawater). Subsequently, the fish are transferred to seawater where they grow into adults. These ontogenic events are likely to impact the microbiome of fish. In the present study, we employed a 16S rRNA gene-based phylotyping technique to assess the transition of the bacterial community at the embryonic stages of Atlantic salmon, and in the intestine of the fish from first feeding stage to the 80-week post-hatch stage. Additionally, we predicted the functional potential of the bacterial communities associated with different life stages of the fish.

## Results

The sequences were clustered into 1442 OTUs, and the rarefied data, with a depth of 2400 sequences/sample, were used to calculate the alpha and beta diversity indices (Additional file 2: Fig. S1a, b). The succession of the ontogeny-associated microbiota of the 4 groups is described in the present study.

### Hatching reflects a shift in the diversity and composition of the microbiota

The bacterial communities of the early developmental stages were examined. The diversity indices (Shannon index and PD Whole tree) of the communities of HL were significantly higher (p<0.05, Fig. 2a) compared with the EE and EBH communities. The evenness associated with the three stages did not significantly vary (p>0.05, Fig. 2a). The community compositions of the early developmental stages were significantly different (Figs. 2b, c; Additional file 6: Table S3a; p<0.01, R>0.6, based on weighted and unweighted UniFrac distances).

**Fig. 1.**
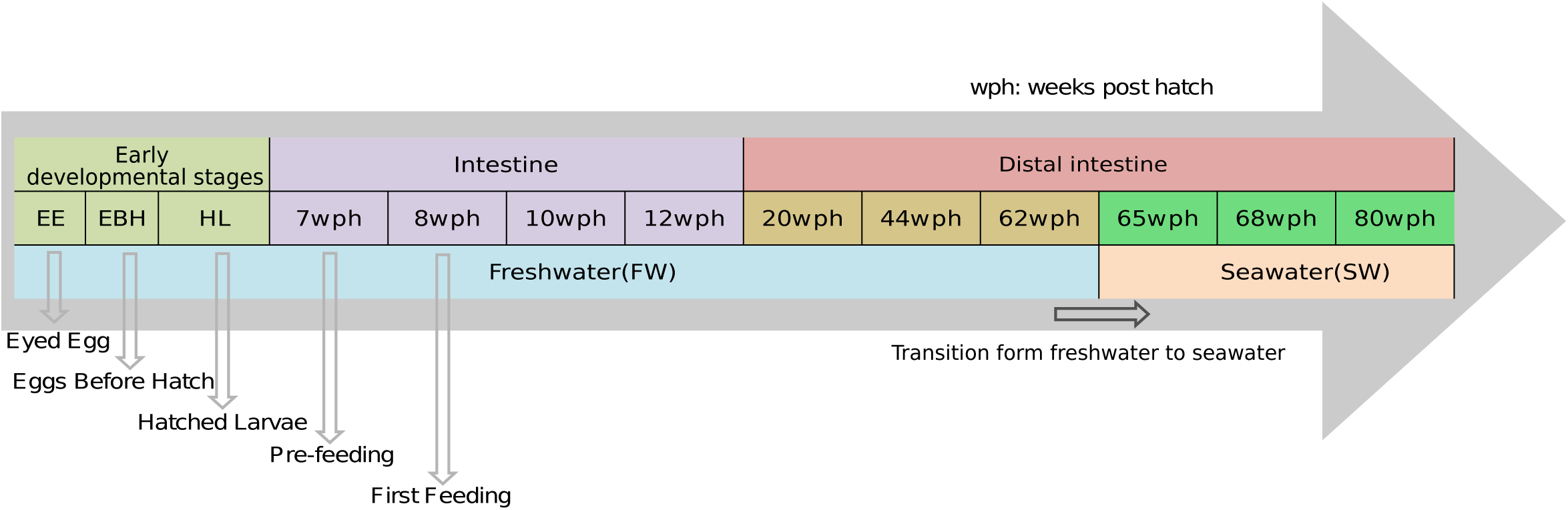
Ontogenetic timeline of salmon depicting the successive developmental stages that were targeted in the present study: early developmental stages; early freshwater stages; late freshwater stages; seawater stages. wph: weeks post hatching.

**Fig. 2.**
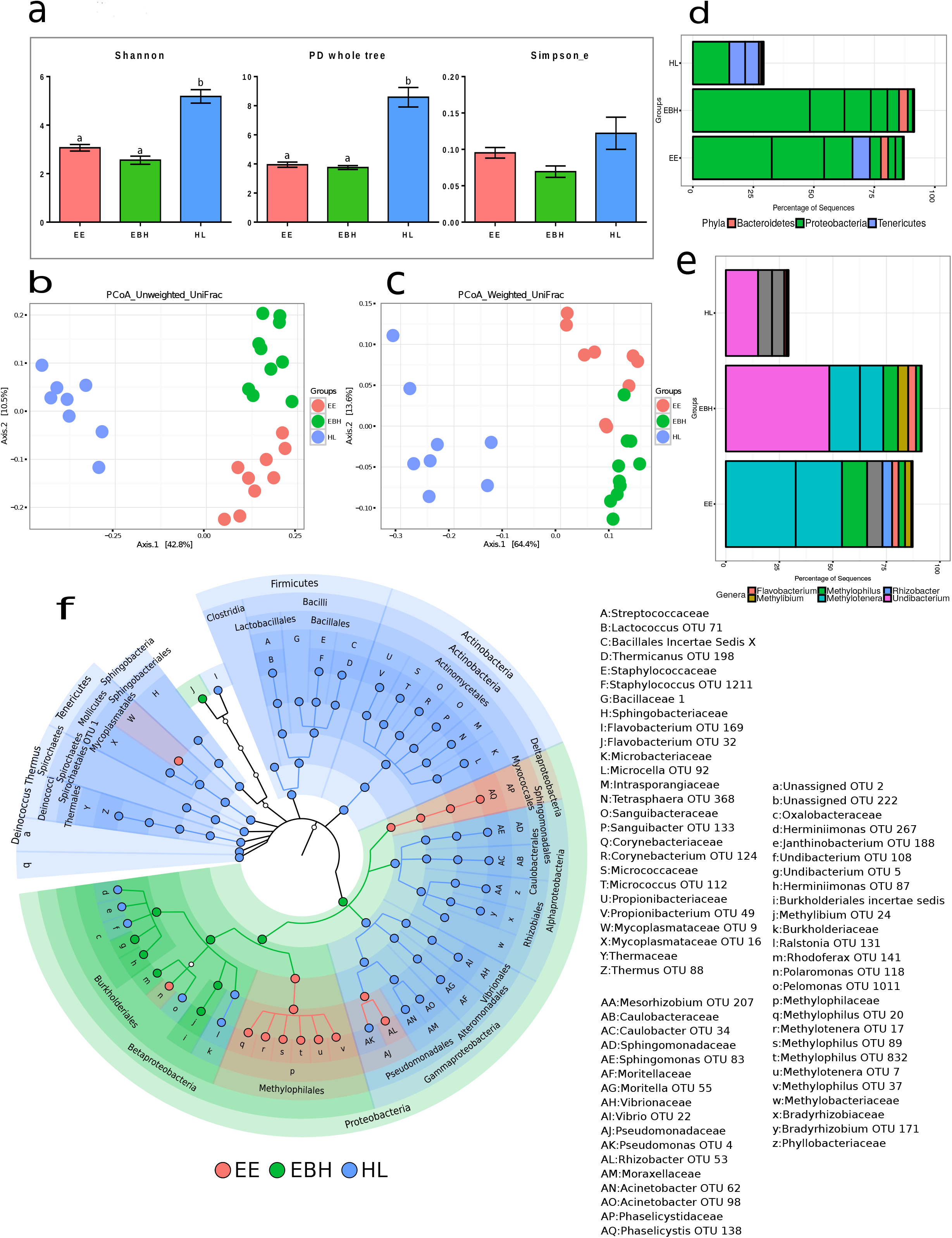
Plots showing the comparisons of microbiota associated with the early developmental stages (EE, EBH, and HL) of Atlantic salmon. Stage-specific colour coding was used for Figures a, b, c, and f. (a) Alpha diversity indices (Shannon index, PD whole tree, and Simpson evenness) of the bacterial communities and (b, c) UniFrac distances-based PCoA. The mean relative abundance of the 10 most abundant OTUs at the (d) phylum and (e) genus levels. The OTUs are coloured according to their taxonomic classification, and OTUs without any assignment are shown in grey. (f) Cladogram showing the significantly abundant taxonomic groups in each of the stages, identified based on the LEfSe (p<0.05 and effect size >3.5).

*Proteobacteria* were significantly abundant in the EBH (Fig. 2f), although the results obtained for the EE (including *Methylotenera* and *Methylophilus*) and HL stages (*Undibacterium*) also showed the most dominant OTUs under this phylum (Fig. 2d, e). The proportion of *Proteobacteria* decreased from the EBH to the HL (Fig. 2d), whereas those of *Actinobacteria, Tenericutes, Firmicutes, Bacteroidetes, Deinococcus-Thermus, Spirochaetes* (also identified as biomarkers, Fig. 2f) increased. *Deltaproteobacteria* in the EE and *Betaproteobacteria* in the EBH were observed as significantly abundant classes under the phylum *Proteobacteria*. The members of the orders *Methylophilales* (*Betaproteobacteria*) and *Myxococcales* (*Deltaproteobacteria*) were abundant in the EE, whereas most of the OTUs under the order *Burkholderiales* were significantly abundant in either the EBH or the HL (Fig. 2f). The OTUs belonging to the orders *Pseudomonadales, Alteromonadales, Virbionales, Rhizobiales, Caulobacterales, Sphingomonadales, Actinomycetales, Bacillales, Lactobacillales, Spingobacteriales, Mycoplasmatales, Spirochaetales*, and *Thermales* were significantly abundant in the HL. Two classes of *Proteobacteria* were significantly abundant in the HL: *Alpha*- and *Gammaproteobacteria*. Furthermore, all OTUs under the above-mentioned classes, except one OTU belonging to the *Rhizobacter*, were significantly abundant in the HL (Fig. 2f). The significantly abundant OTUs under each phylum and their effect sizes are listed in Additional file 7: Table S4a.

### Successional changes in the diversity and composition of the intestinal bacterial community of fish at the early freshwater stages

The alpha diversity indices of the communities associated with the intestine of fish at the early freshwater stages did not significantly vary (Fig. 3a). The intestinal bacterial communities of the fish at the early freshwater stages were significantly different (Fig. 3c; Additional file 6: Table S3b; p<0.01, R>0.5; based on weighted UniFrac distances 7, 8, 10 vs. 12 wph).

**Fig. 3.**
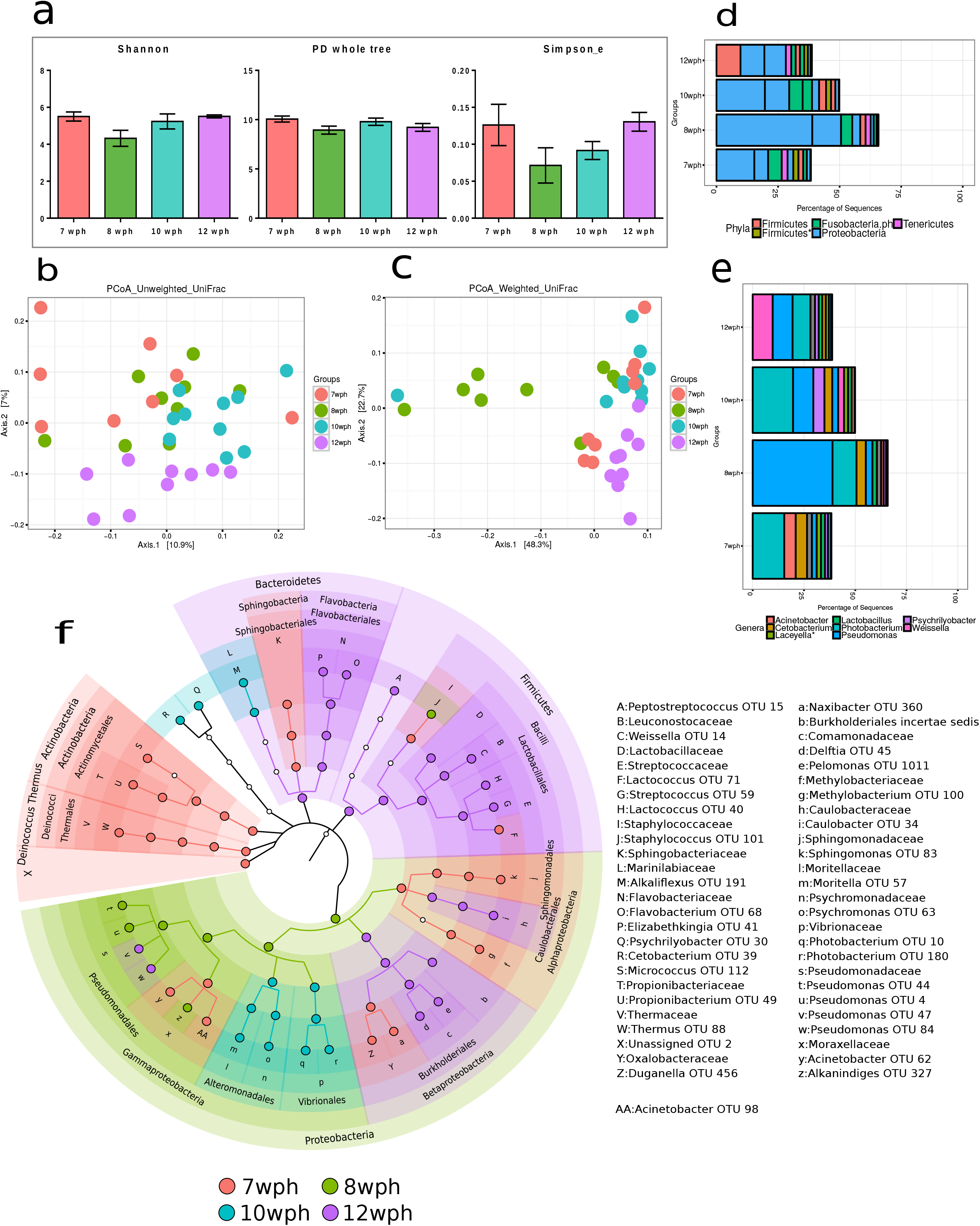
Plots showing the comparisons of the microbiota associated with the whole intestine of Atlantic salmon in freshwater (7, 8, 10 and 12 wph). Stage-specific colour coding is used for Figures a, b, c, and f. (a) Alpha diversity indices (Shannon index, PD whole tree, and Simpson evenness) of the bacterial communities and (b, c) UniFrac distances-based PCoA. The mean relative abundance of the 10 most abundant OTUs at the (d) phylum and (e) genus levels. OTUs are coloured according to their taxonomic classification, and the OTUs without any assignment are shown in grey. (f) Cladogram showing the significantly abundant taxonomic groups in each of the stages, identified based on the LEfSe (p<0.05 and effect size >3.5).

*Proteobacteria* was the dominant phylum in all the stages (Fig. 3d). However, as the fish were growing the changes were evident from the significantly abundant OTUs associated with the stages (Fig. 3f). The phylum *Proteobacteria* was significantly abundant at 8 wph, primarily reflecting the abundance of the OTUs of the order *Pseudomonadales*, whereas *Vibrionales, Alteromonadales* and the families and genera under these orders were significantly abundant at 10 wph. The significantly abundant OTUs belonging to *Comamonadaceae* under *Burkholderiales* made *Betaproteobacteria* a significant feature at 12 wph, whereas the OTUs of *Oxalobacteriaceae*, belonging to *Betaproteobacteria*, were significantly abundant at 7 wph. *Alphaproteobacteria* was significantly abundant at 7 wph, comprising the OTUs belonging to *Sphingomonadales* and *Methylobacteriaceae*. However, *Caulobacteriales* (*Alphaproteobacteria*) and its members were significantly abundant at 12 wph (Fig. 3f). The phyla *Actinobacteria* and *Deinococcus-Thermus* were significantly abundant at 7 wph (Fig. 3f). *Bacteroidetes* was significantly abundant at 12 wph, primarily reflecting the significant abundances of the *Flavobacterial* lineage, whereas the class *Sphingobacteria* (*Bacteroidetes*) was significantly abundant at 7 wph. *Firmicutes* and most of the members of this phylum, particularly the OTUs belonging to the class *Bacilli*, were significantly abundant at 12 wph. Additional file 7: Table S4b lists the significantly different OTUs and their effect sizes under each phylum.

### Successional changes in the diversity and composition of the distal intestinal community of fish at the late freshwater stages

The Shannon index of the bacterial communities at 20 and 44 wph were significantly different (Fig. 4a). However, the richness (PD whole tree) and evenness (Simpson’s evenness) of the communities, when considered individually, did not significantly vary (Fig. 4a). The fish at the late freshwater stages had significantly different [Fig. 4b, c; Additional file 6: Table S3c; p*<*0.01, R*>*0.8, based on unweighted (20 vs. 44 wph) and weighted UniFrac distances (20 vs. 44, 62 wph)] bacterial communities. *Firmicutes* was the most dominant phylum in the distal intestine at 20, 44 and 62 wph (Fig. 4d). In addition, 2 OTUs with taxonomy prediction confidence estimates *<*0.5 (hence excluded from the LEfSe analysis) belonging to the phylum *Firmicutes (*indicated using *, Fig. 4d; including the genus *Laceyella*, Fig. 4e) were also predominant in this group of fish. The phylum *Firmicutes* and the OTUs under this group, *Lactobacillales* and *Bacillales*, comprising the class *Bacilli*, were significantly abundant at 20 wph (Fig. 4f). The class *Clostridia*, however, was significantly abundant at 62 wph (primarily reflecting one OTU belonging to *Anaerofilum)*. Other OTUs belonging to *Peptostreptococcaceae* and some unassigned OTUs under *Clostridiales* were significantly abundant at 44 wph (Fig. 4f). While the phylum *Tenericutes* and its members were significantly abundant at 62 wph, the phylum *Bacteroidetes* and its members were significantly abundant at 20 wph. At the phylum level, *Proteobacteria* was not a significant feature of any of the stages. However, the classes under this group *(Alpha-, Beta-* and *Gammaproteobacteria)* were significant features at 20 wph (Fig. 4f). Interestingly, at the order level, the significantly abundant features belonged to different stages, including *Rhizobiales* (20 wph) and *Caulobacteriales* (62 wph) of *Alphaproteobacteria, Pseudomonadales* (20 wph*), Enterobacteriales* (20 wph*), Vibrionales* (44 wph) and *Aeromonadales* (62 wph) of *Gammaproteobacteria* (Fig. 4f). Additional file 7: Table S4c lists the significantly different OTUs and their effect sizes under each phylum.

**Fig. 4.**
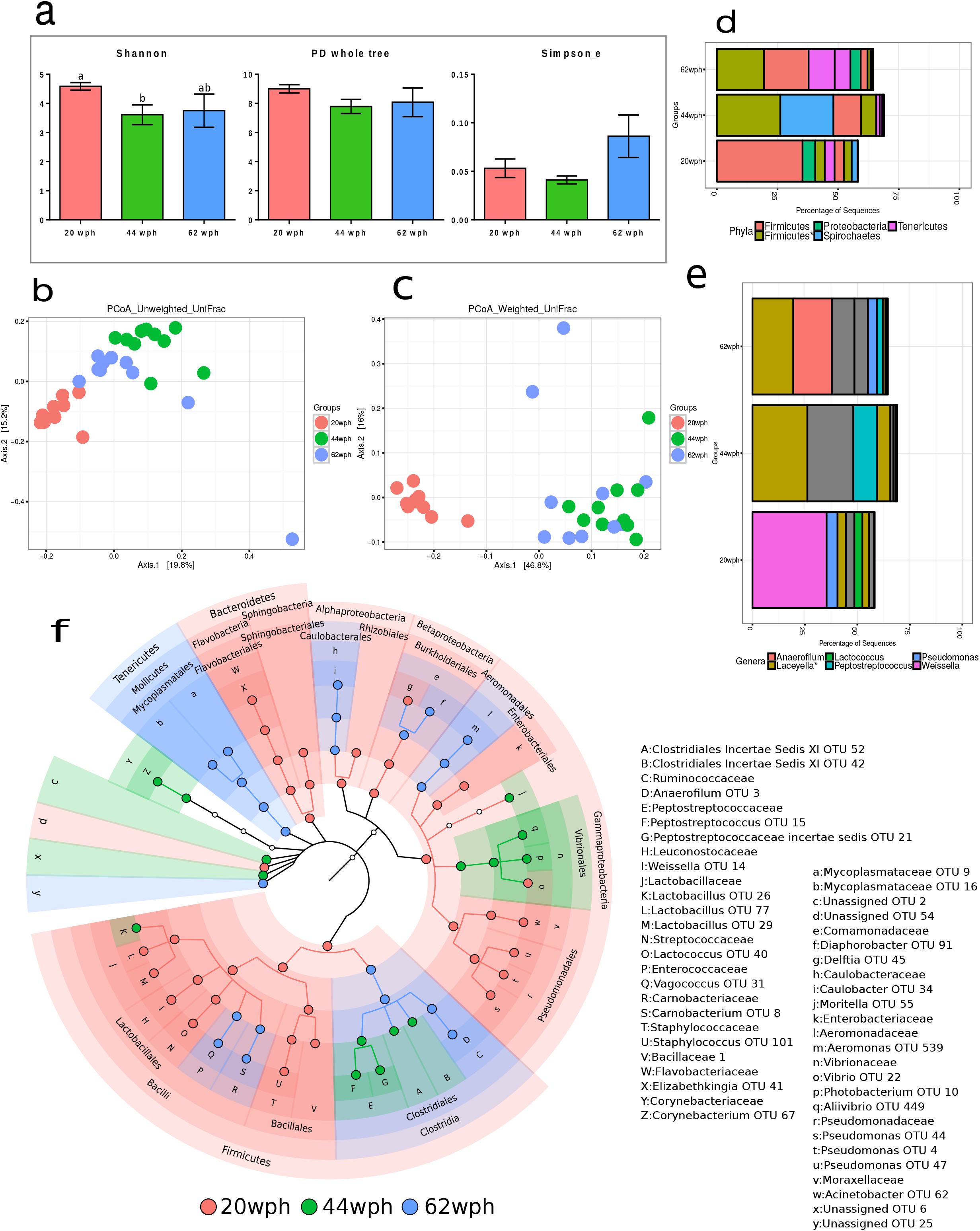
Plots showing the comparison of the microbiota associated with the distal intestine of Atlantic salmon in freshwater (20, 44 and 62 wph). Stage-specific colour coding is used for Figures a, b, c, and f. (a) Alpha diversity indices (Shannon index, PD whole tree, and Simpson evenness) of the bacterial communities and (b, c) UniFrac distances-based PCoA. (d) The mean relative abundance of the 10 most abundant OTUs at the (d) phylum and (e) genus levels. OTUs are coloured according to their taxonomic classification, and the OTUs without any assignment are shown in grey. (f) Cladogram showing the significantly abundant taxonomic groups in each of the stages, identified based on the LEfSe (p<0.05 and effect size >3.5).

### Successional changes in the diversity and composition of the distal intestinal community of seawater fish

The Shannon indices of the communities associated with the distal intestine of the Atlantic salmon in seawater (65, 68 and 80 wph stages) were significantly different (Shannon index; Fig. 5a, p<0.05). The bacterial community compositions of fish at the seawater stages were significantly different (Fig. 5b, c; Additional file 6: Table S3d; p<0.01, R<0.6, based on unweighted and weighted UniFrac distances). *Firmicutes\** was the dominant phylum in the distal intestine of the fish in seawater, particularly at 65 and 80 wph (Fig. 5d). The 2 OTUs (with low taxonomic assignment confidence, <0.5) belonging to the genus *Laceyella* (phylum *Firmicutes*) were also predominant at 65, 68 and 80 wph (Fig. 5e). The phylum *Spirochaetes* was also predominant in the distal intestine at 68 wph. *Actinobacteria, Tenericutes* and *Firmicutes* were the significantly abundant phyla at 65 wph. *Spirochaetes* and *Bacteroidetes* were the significant phyla at 68 and 80 wph, respectively. Under *Firmicutes*, one OTU belonging to *Weissella* was a feature of the 80 wph, making *Lactobacillales* a significant feature at 80 wph. Although at 65 wph more significantly abundant taxonomic biomarkers were observed for the phylum *Proteobacteria*, phylum-level significant abundance was not detected. The classes *Alphaproteobacteria, Epsilonproteobacteria* and their members were significantly abundant at 65 wph. Under *Proteobacteria*, the orders *Alteromonadales, Pseudomonadales* and 2 OTUs belonging to the genus *Vibrio* were significantly abundant at 65 wph (Fig. 5f). Under *Pseudomonadales*, 2 OTUs of *Psychrobacter* and *Pseudomonas* were the significantly abundant features at 80 wph (Fig. 5f). Additional file 7: Table S4d lists the significantly different OTUs and their effect sizes under each phylum.

**Fig. 5.**
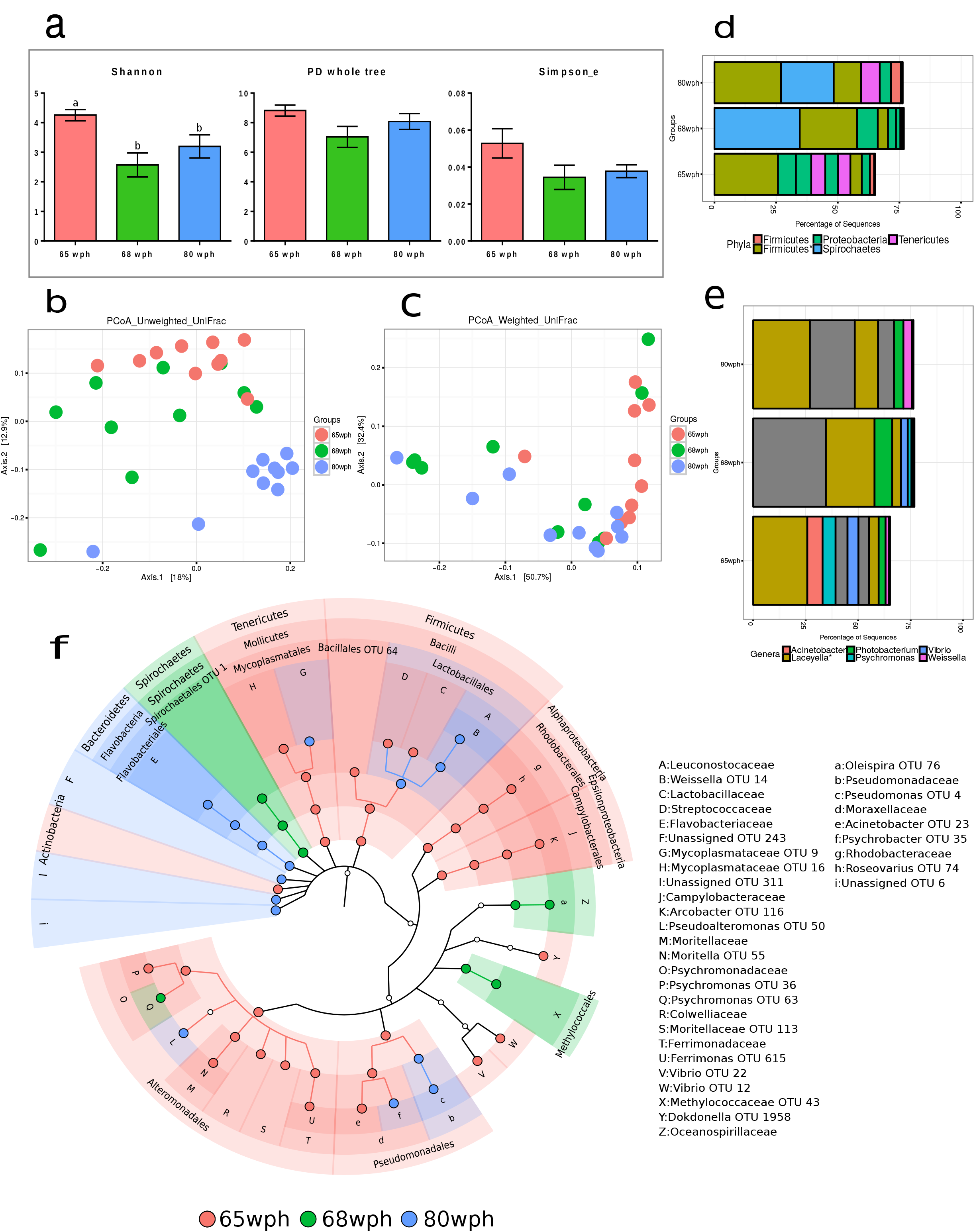
Plots showing the comparison of the microbiota associated with the distal intestine of Atlantic salmon in seawater (65, 68 and 80 wph). Stage-specific colour coding is used for Figures a, b, c, and f. (a) Alpha diversity indices (Shannon index, PD whole tree, and Simpson evenness) of the bacterial communities and (b, c) UniFrac distances-based PCoA. (d) The mean relative abundance of the 10 most abundant OTUs at the (d) phylum and (e) genus levels. OTUs are coloured according to their taxonomic classification, and the OTUs without any assignment are shown in grey. (f) Cladogram showing the significantly abundant taxonomic groups in each of the stages, identified based on the LEfSe (p<0.05 and effect size >3.5).

The bacterial compositional shift at the phylum-level along all the stages sampled is shown in Fig. 6. The prominence of *Proteobacteria* decreased slowly, and when the fish was in seawater *Firmicutes* and *Spirochaetes* surpassed *Proteobacteria* in dominance.

**Fig. 6.**
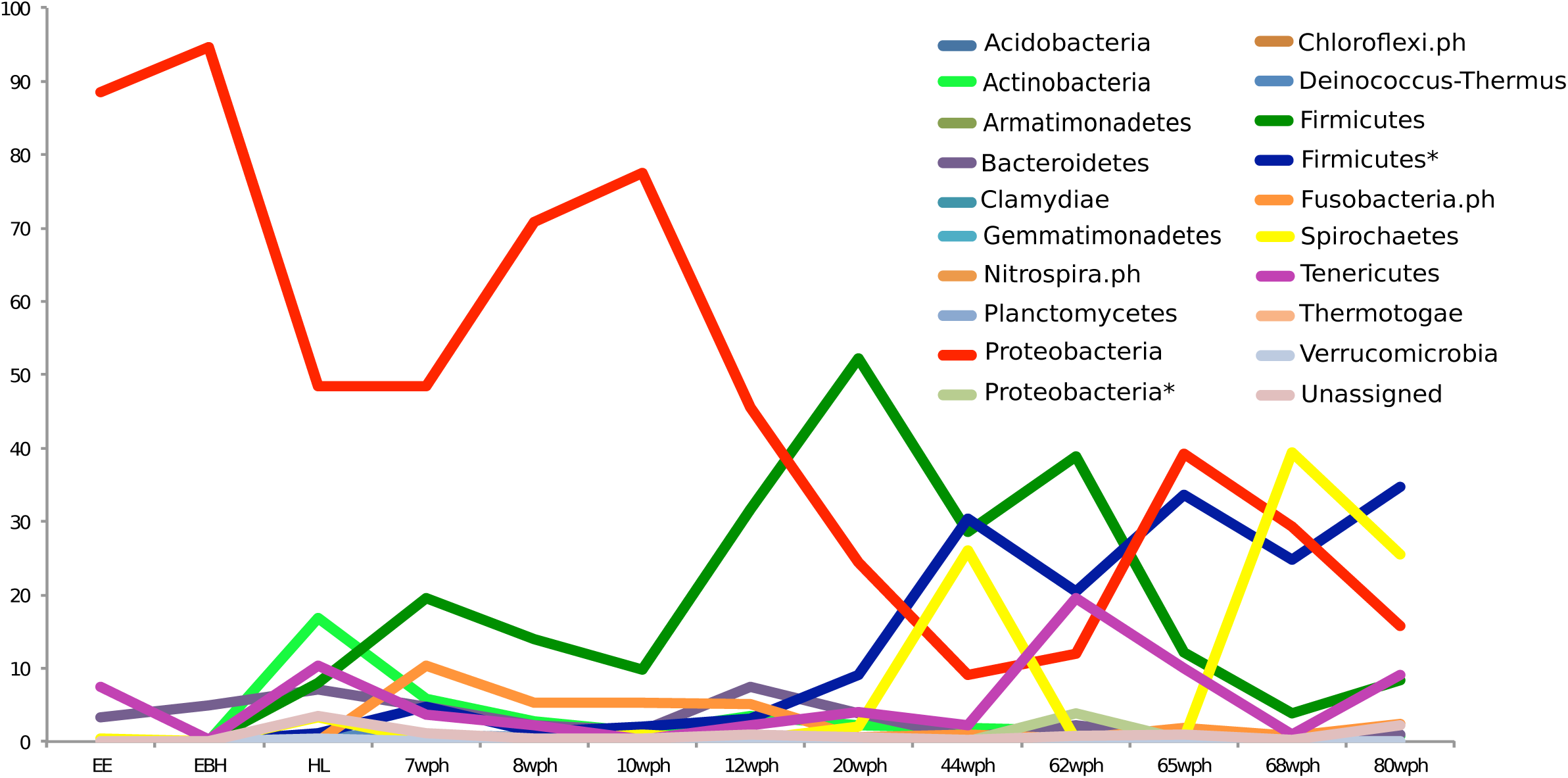
Overview of the phylum-level shifts in the bacterial communities at the different life stages of Atlantic salmon originating from a single cohort. * indicates that phyla with taxonomy assignment confidence below 0.5.

### Comparison of the communities associated with the hatchlings and the intestine of fish at the early freshwater stage

The alpha diversity indices of the communities at the HL and 7 wph stages did not significantly vary (Additional file 2: Fig. S2a). However, the bacterial community compositions at the HL and 7 wph stages were significantly different (Additional file 2: Fig. S2c; Additional file 6: Table S3e – p<0.01, R>0.6; based on weighted UniFrac distances).

The dominant OTUs of the 2 stages (Additional file 2: Fig. S2d, e) and their phylum level biomarkers (Additional file 2: Fig. S2f) indicate differences in the bacterial communities. *Actinobacteria, Tenericutes* and *Spirochaetes*, and their members were significantly abundant in the HL, whereas, *Fusobacteria*.ph and *Firmicutes*, and their members were significant features at 7 wph (Additional file 2: Fig. S2f). *Proteobacteria* was not identified as a biomarker, but the classes *Beta-* and *Gammaproteobacteria* and the associated OTUs were significantly abundant in the HL and at 7 wph. The effect sizes of the respective features are provided in Additional file 7: Table S4e.

### Comparison of the communities associated with the whole and distal intestine of fish at the freshwater stage

The Shannon index and evenness of the bacterial communities at 20 wph were significantly lower (p<0.05) compared to those at 12 wph (Additional file 3: Fig. S3a). However, the richness (PD whole tree) associated with the two stages was similar. The bacterial community compositions of the two stages were significantly different (Additional file 3: Fig. S3c; Additional file 6: Table S3e; p<0.01, R>0.8, based on the weighted UniFrac distances).

*Firmicutes* and *Proteobacteria* were the dominant phyla in the two stages examined (Additional file 3: Fig. S3d). The phylum *Actinobacteria, Fusobacteria*.ph, *Bacteroidetes* and *Proteobacteria* were significantly abundant at 12 wph (Additional file 3: Fig. S3f), whereas *Tenericutes, Spirochaetes* and *Firmicutes* were significantly abundant at 20 wph. Under *Firmicutes*, the class *Clostridia* and its members were significantly abundant at 12 wph, whereas *Bacilli* were significantly abundant at 20 wph (Additional file3: Fig. S3f). Under *Bacilli*, 4 OTUs belonging to *Lactobacillus, Streptococcus, Vagococcus* and *Filibacter* were the significant features at 12 wph. The effect sizes of the respective features are provided in Additional file 7: Table S4f.

### Comparison of the communities associated with the distal intestine of freshwater and seawater fish

There were no significant differences in the diversity indices (Additional file 3: Figs. S4a, b, c; Additional file 6: Table S3e; p<0.01, R<0.6) associated with 62 and 65 wph. *Firmicutes, Tenericutes* and *Proteobacteria* were the dominant phyla at the two stages examined (Additional file3: Fig. S4d). *Bacteroidetes* and *Firmicutes* were abundant at 62 wph (freshwater), whereas *Proteobacteria* was significantly abundant at 65 wph (seawater) (Additional file 3: Fig. S4f). Some members of *Proteobacteria* namely, *Caulobacterales, Burkholderiales* and *Pseudomonadaceae* were the abundant features at 62 wph (Additional file 3: Fig. S4f). The OTUs under *Firmicutes*, including *Clostridiales, Bacillales, Streptococcus* and *Leuconostocaceae*, were the significant features at 65 wph. The effect sizes of the respective features are provided in Additional file 7: Table S4g.

### Presumptive functions of the communities at different stages

The presumptive functional pathways associated with the microbiota at different stages were analysed to identify the stage-specific significant functional potential of these bacteria. The NSTI (Nearest Sequenced Taxon Index) scores (Langille *et al*., 2013) corresponding to each of the predictions are provided in Additional file3: Fig. S5. Additional file 8: Table S5a lists the five most abundant KEGG modules and differentially abundant features (p<0.01 and effect size >0.75) at each stage. The functional potential of the community of the HL was significantly different from those of the EE and EBH (Fig. 7, Additional file 9: Table S6a; *p*<0.01, R>0.85). The functions associated with the communities of the fish at the early freshwater and the seawater stages were not significantly different (Fig. 7, Additional file 9: Table S6b, d). The functional potential of the distal intestinal community of the fish at the early freshwater stages (20 wph) was significantly different from that of the fish at the late freshwater stages (44 and 62 wph, Fig. 7, Additional file 9: Table S6c; p<0.01, R>0.6).

**Fig. 7.**
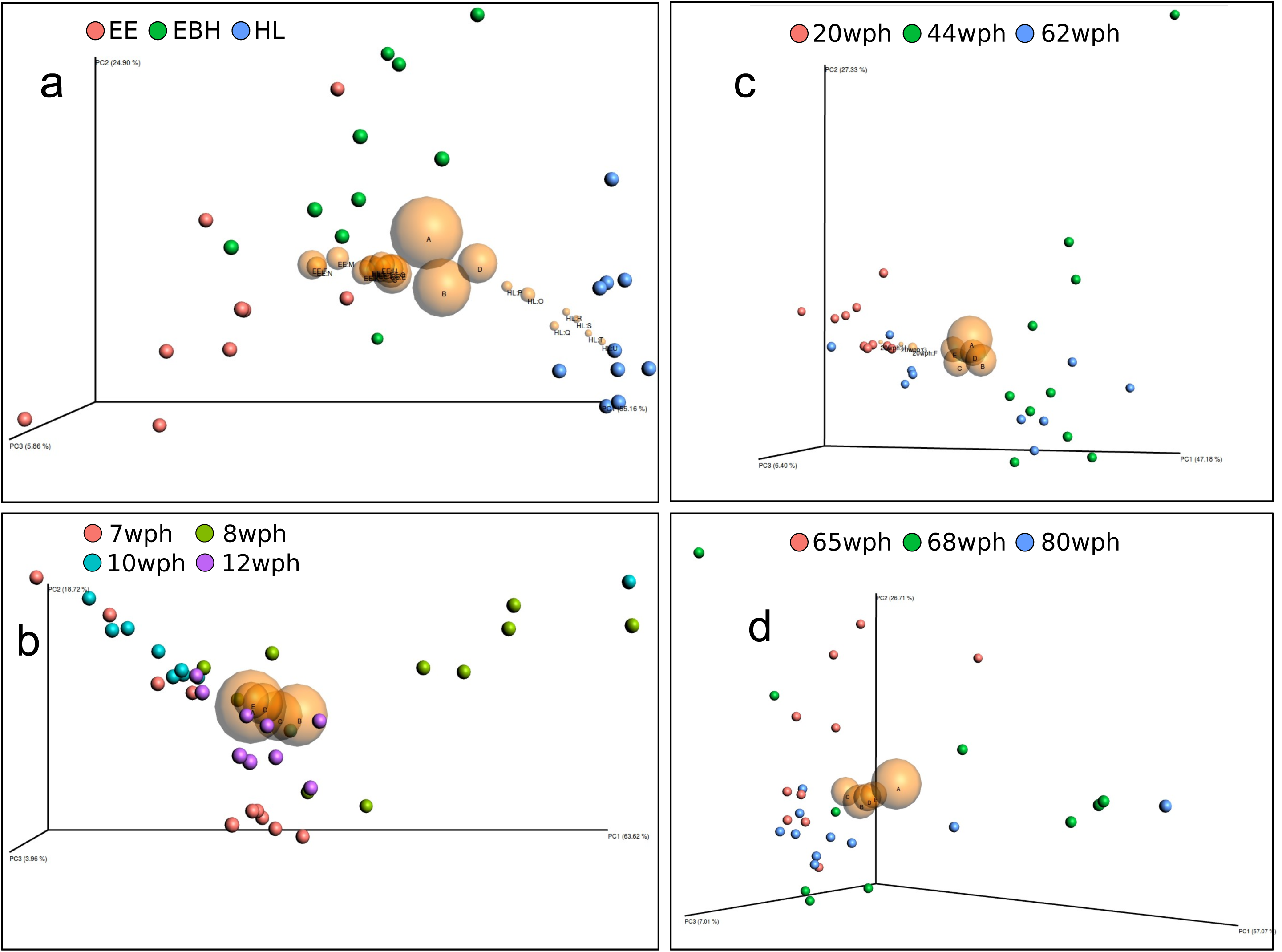
Comparison of the functional pathways associated with different life stages. Bray-Curtis dissimilarities based on the abundances of different functional pathways are shown using PCoA biplot. Stage-specific colour coding is used for the figures. (a) Early developmental stages, (b) whole intestine, (c) distal intestine (freshwater) and (d) distal intestine (seawater). The seven most abundant features of each stage (with an average abundance >2.5%) are shown using yellow spheres. The size of the spheres is indicative of the relative abundances of the features and is named alphabetically (A to E, in the order of decreasing abundance; see Additional file 8: Table S5a). Significantly abundant features (p<0.05 and effect size >0.75) belonging to the respective stages are also represented using yellow spheres, labelled alphabetically from F, in order of decreasing abundance. The names of the features are provided in Additional file 8: Table S5a.

The seven pathways that were significantly abundant across stages were branched-chain amino acid transport system, peptides/nickel transport system, riboflavin biosynthesis, multiple sugar transport system, pentose phosphate pathway, phosphate transport system and glycolysis.

## Discussion

The present study profiled the bacterial communities of Atlantic salmon to examine the progressive transition of these fish communities during the early embryonic stages (EE, EBH and HL), in the intestine during the early freshwater stages (7, 8, 10, 12 wph), in the distal intestine of the late freshwater stages (20, 44, 62 wph), and in the distal intestine of the seawater stages (65, 68 and 80 wph). Shifts in the predicted functional content of the communities are also discussed. The gut microbiota of the aquacultured species (grass carp, *Ctenopharyngodon idella;* Chinese perch, *Siniperca chuatsi;* and southern catfish, *Silurus meridionalis*) from the same regional pool are reported to be similar as well as developmental stage-dependent, and they are distinct when compared with the bacteria in water [14]. It is plausible that the similar deterministic processes also regulate the succession in the bacterial communities of the Atlantic salmon.

Fish eggs are colonized by diverse microbial communities [15, 16]. In the present study, the bacterial community associated with the whole organism was examined up to the hatching stage. The alpha diversity indices at the embryonic stages (egg surface) were the lowest compared with the hatched larvae. The predominant OTUs associated with the embryonic stages of Atlantic salmon belonged to *Methylotenera* and *Undibacterium. Vibrio fischeri* and *Leucothrix mucor* were abundant on cod eggs, whereas *Moraxella* and *Alcaligens* were abundant on halibut eggs. In addition, microbiota of cod larvae was highly distinct from those of their environment and live feed [9, 15, 16]. Taken together, these findings suggest that the early life communities are species- and stage-specific.

The transition from eyed eggs (EE) to those prior to hatching (EBH) was characterised based on changes, particularly at the genus level: *Methylotenera* and *Methylophilus* were dominant in the EE, whereas *Undibacterium* was dominant in the EBH and HL. These communities are likely to be egg surface-specific [17, 16, 18], and the mechanisms causing such shifts are not clear yet, although neutral and non-neutral assembly models have been proposed for zebrafish [19]. As zebrafish ages, the assembly of the associated bacterial community is not decided according to chance and dispersal, but through microbial interactions, active dispersal, or host selection [19]. The hatchling-associated community was significantly diverse (phylogenetically different) compared with the communities prior to hatching. Hatching is a critical process because the sterile embryo contacts the microbe-rich environment [16, 20] when the immune system of the organism is still immature [21]. These diverse community members might aid the host in defence against pathogens [22, 16, 23]. From the hatching stage onward, major mucosal organs, such as the gills, skin and gut, become functionally active [24], and the specific phylotypes that colonize these microenvironments might play key roles in the normal development of these organs [25-27, 23, 16]. In addition, at this stage, oxygen uptake changes from cutaneous to pharyngeal [28], and this development could affect the community composition. These ontogenic changes might contribute to the HL-associated diverse bacterial community.

After the formation of the gut, i.e., 7 weeks after hatching, the bacterial community associated with the whole intestine was assessed. The alpha diversity indices of the intestinal microbiota at 7 wph (prior to first feeding) and the stages after feeding (8, 10 and 12 wph) did not significantly vary. Feeding led to a transition of the rainbow trout larval intestine from a *Bacteroidetes-dominant* to a *Firmicutes-* and *Proteobacteria-dominant* community [27]. The observations in the present study suggest that feeding causes a phylum-level shift to *Proteobacteria* (at 8 wph) and *Bacteroidetes* (as a result of the *Flavobacterial* lineage, at 12wph), and *Firmicutes* (primarily reflecting the genus *Weissella*, at 12 wph).

The distal intestine was clearly distinguishable at 20 wph; therefore, the bacterial community associated with this intestinal region was analysed from this time point. The significant decrease in the alpha diversity index (20 vs. 44 wph) could reflect the less diverse community at 44 wph and the overrepresentation of *Spirochaetes* in the distal intestinal microbiota associated with this stage. Similar to the findings in the present study, *Spirochaetes* are highly abundant in other carnivorous fish, including mahi-mahi (*Coryphaena hippurus*) and great barracuda (*Sphyraena barracuda*) [29]. In the present study, the phylum *Firmicutes* was significantly abundant at 20 wph. The genera *Weissella, Laceyella\** and *Anaerofilum* were the predominant contributors to the significant abundance of *Firmicutes*. Rainbow trout, also presents a high abundance of *Firmicutes*, with OTUs belonging to Bacilli as the predominant type [30]. This observation is similar to the findings in the present study. In contrast, members of *Bacilli* were not abundant in the gut of the cyprinids common carp (*Cyprinus carpio*) and zebrafish (*Danio rerio*) [31, 32], indicating the importance of *Firmicutes* in salmonids. Furthermore, the phylum *Tenericutes* became significantly abundant just prior to the transfer of these fish to seawater. *Tenericutes* are highly abundant in salmon (both in freshwater and seawater) [33, 13] and trout intestines [34].

*Firmicutes* were significantly abundant soon after the fish were transferred to seawater, and the OTUs belonging to *Laceyella*\* remained predominant. In addition, the OTUs belonging to *Spirochaetes, Proteobacteria* and *Tenericutes* were also prominent. The dominance of *Spirochaetes* at 44 wph and the significant abundance of the phylum at 68 wph suggest an important role of this taxon in the gut microbiota of carnivorous fish. During the seawater stages, the alpha diversity index of the distal intestinal community significantly decreased with time. The lower alpha diversity indices and the overabundance of the few phylotypes in the microbiota of the intestine [13] and skin [35] of adult Atlantic salmon and rainbow trout intestine [34] have been previously documented. Changes in the phylum *Tenericutes* (mainly *Mycoplasma* spp.) during development were minimal in the present study. Although *Tenericutes* were part of the microbiota at the early developmental stages and were significantly abundant in the HL and the distal intestine at 62 and 65 wph, the proportion of this phylum (20% at 62 wph) was much less compared with the study by Holben *et al*. [33], who reported 70-90% *Tenericutes* in most of their samples. Another study on the transition in the community composition of the wild Atlantic salmon by Llewellyn *et al*. [16] showed that the proportion of *Mycoplasma* spp. increased consistently with development and it was most abundant in the seawater fish. In addition, previous reports on the abundance of *Mycoplasma* spp. in the intestine are contrasting; Llewellyn *et al*. [16] and Holben *et al*. [33] found an over dominance, whereas Zarkasi *et al*. [36, 37] detected only sporadic occurrence of the species. These discrepancies could be because of the genetic background or the geographical locations of the fish sampled.

We also examined the diversity and significantly abundant phyla associated with the whole animal, whole intestine, and distal intestine of the fish in freshwater and seawater by conducting the following comparisons; HL vs. 7, 12 vs. 20, and 62 vs. 65 wph. These comparisons revealed significant differences in the diversity indices and the composition of the communities, which was even reflected at the phylum level. *Firmicutes* were significantly abundant in the whole and distal intestine of the fish in freshwater but not in the distal intestine of the fish in seawater. *Proteobacteria*, however, were significantly abundant in the whole and distal intestine of the fish in seawater. These results indicate that the *Proteobacteria*-rich community in the early intestine changes to a *Firmicutes*-rich distal intestinal community in freshwater. However, when the fish were introduced into seawater, *Proteobacteria* regained significant abundance. This transition to a *Proteobacteria*-rich community when the fish enters seawater has been previously recorded in fish skin microbiota [35, 38]. A meta-analysis also revealed the differences in the gut bacterial community compositions of freshwater and the seawater fishes [39].

### The presumptive functional pathways of the bacterial c<u>omm</u>unities of Atlantic salmon

The mean weighted NSTI scores for the communities at different stages ranged from 0.43 ± 0.006 to 0.295 ± 0.037. In general, the early stages had lower NSTI values compared with the stages from 44 wph onward. This finding indicates that the metagenomes of the communities associated with the distal intestine were predicted based on higher taxonomic levels, which make these data less accurate. The five core functions of the bacteria associated with Atlantic salmon included biosynthetic (riboflavin) and transport pathways. These functions were associated with all stages of development and did not vary, despite differences in the community composition, indicating their importance throughout development. The core metabolic functional potential of bacteria can be similar, even when there are differences in the phylogenetic content [40]. However, the functional pathways that were significantly represented in the EE and HL and in the intestine at 20 wph indicate the specific needs of the associated bacteria or host. These pathways included biosynthesis of pantothenate, biotin, ADP-L-glycero-D-manno-heptose, heme, methionine and ketone body. The significance of these functional pathways in relation to the physiological needs of the fish should be explored further.

### Conclusion

The present study examined the transition of the embryonic and intestinal bacterial communities of Atlantic salmon. Stage-specific microbial signatures were evident at the phylum level. *Proteobacteria* was the most abundant phylum in eggs, and its abundance decreased in the hatchlings. The diversity of the hatchling-associated community increased, reflecting the significant abundance of *Actinobacteria, Firmicutes, Tenericutes, Spirochaetes* and *Deinococcus-Thermus*. In the intestine of the fish at the early freshwater stages, the phylum *Proteobacteria* was dominant. Although *Firmicutes* and *Bacteroidetes* subsequently became the significantly abundant phyla, only the dominance of *Firmicutes* was evident in the distal intestine of the fish at the late freshwater stages. After the fish were in seawater, *Proteobacteria* again became the significantly abundant phylum. However, *Firmicutes, Spirochaetes, Tenericutes* and *Actinobacteria* were the significantly abundant phyla as the fish adapted to its life in seawater. The functional redundancy of the taxonomically dissimilar communities associated with the different stages are likely related to the specific needs of the associated bacterial communities or host.

## Methods

### Biological material

Samples (n=10) from selected life stages (up to smolts) of the fish were procured from a local hatchery (Cermaq AS, Hopen, Bodø, Norway). The smolts were transported to the research station at Nord University and further reared in a seawater facility at the station. More information is provided in Additional Methods.

### Sampling

The fish were euthanized prior to sampling. The samples from the successive developmental stages were classified in 4 groups as follows: i) the whole organism (early developmental stages: eyed egg stage, EE; embryo before hatching, EBH; and hatched larvae, HL); ii) the whole intestine of the fish at early freshwater stages (7, 8, 10 and 12 weeks post hatch, wph); iii) the distal intestine of fish at the late freshwater stage (20, 44 and 62 wph); and iv) the distal intestine of fish at the seawater stage (65, 68 and 80 wph) (Fig. 1). Further details are provided in Additional file 1: Methods and Additional file 4: Table S1.

### DNA extraction, preparation of the sequencing libraries (V3-V4 region), library quantification and sequencing

DNA from the samples was extracted using the QIAamp Fast DNA Stool Mini Kit (Qiagen, Nydalen, Sweden). The samples were processed according to the manufacturer’s protocol, with few modifications as detailed in the Additional file 1: Methods.

A paired end, dual index protocol was adopted to amplify and prepare the 16S rRNA gene (V3-V4 regions) sequencing libraries [41]. The PCR reactions were performed in a 25 μl reaction volume containing 12.5 μl of Kapa HiFi Hot Start PCR Ready Mix (KAPA Biosystems, Woburn, USA), 2.5 μl of each forward and reverse primer (300 nM), and 7.5 μl of DNA and water. The thermocycling conditions included initial denaturation at 95°C, followed by 35 cycles of 98°C-30s, 58°C-30s and 72°C. The final extension was performed at 72°C for 2min. The PCR products (sequencing libraries) were run on agarose gel and purified, and the libraries were quantified and pooled at equimolar (2 nM) concentrations prior to sequencing (see Additional file 1: Methods).

### Data analysis

UPARSE [42] was used for quality filtering and OTU clustering. Forward reads comprising the V3 region (see Additional file 1: Methods) of the 16S rRNA gene were quality filtered, truncated to 200 bp, dereplicated, and abundance sorted, and reads with less than 10 sequences were discarded. OTUs were clustered at a 97% similarity level, and chimeric sequences were removed using UCHIME [43]. The reads were subsequently mapped to OTUs after searching the reads as a query against the OTU representative sequences. Taxonomic ranks were assigned to the OTUs using the UTAX algorithm (http://www.drive5.com/usearch/manual/utax_algo.html). OTU tables were prepared and split into the 4 study groups (as described in the section Sampling), and comparisons of the bacterial communities in the 4 groups (whole organism, EE, EBH and HL; whole intestine at 7, 8,10 and 12 wph; freshwater distal intestine at 20, 44 and 62 wph; and seawater distal intestine at 65, 68 and 80 wph) were performed separately. To explore the intergroup changes in the diversity and abundances of the associated microbiota, we conducted three additional comparisons: whole organism vs. intestine (HL and 7 wph), intestine vs. distal intestine (12 and 20 wph), and freshwater distal intestine vs. seawater distal intestine (62 and 65 wph). The read statistics of the sequences are provided in Additional file 5: Table S2. For each of the 4 groups the diversity indices were calculated, and the differential abundance analyses were performed separately on the 4 groups using QIIME [44] and LEfSe [45], respectively. The PCoA plot and cladogram showing the differential abundances were created using phyloseq [46] and GraPhlAn [47], respectively. Presumptive metabolic potential was computed using PICRUSt [48], and the resulting gene abundance data were profiled into metabolic pathways using HUMAnN [49], with the default settings. Subsequently, the KEGG modules were analysed using STAMP [50], and the Bray-Curtis dissimilarities were plotted using phyloseq (see Additional file 1: Methods).

## List of abbreviations

16S rRNA: 16S ribosomal RNA
EBH: embryo before hatching
EE: eyed egg stage
FOTS: Forsøksdyrforvatningen tilsyns-og søknadssystem
HL: hatched larvae
HUMAnN: HMP unified metabolic analysis network
KEGG: Kyoto encyclopedia of genes and genomes
LEfSe: linear discriminant analysis effect size
NSTI: nearest sequenced taxon index
OTU: operational taxonomic unit
PCoA: principal coordinate analysis
PCR: Polymerase chain reaction
PD Whole tree: phylogenetic diversity whole tree
PICRUSt: Phylotypic investigation of communities by reconstruction of unobserved states
QIIME: Quantitative insights into microbial ecology
V3 and V4 regions: hypervariable regions 3 and 4
wph: weeks post hatch

## Declarations

### Ethics approval

This study was conducted according to the guidelines of the Norwegian Food Safety Authority (FOTS ID: 7899).

## Availability of data and material

Sample metadata, read statistics, statistical analyses results and additional methods and figures are provided as Additional files. Please contact author for further data request.

## Author contributions

JL and VK conceived the study. JL performed the experiments and data analysis, wrote and redressed the manuscript. VK scrutinized the data, read and redressed the manuscript. DS, JF and TM discussed the experimental design, read and critically edited the manuscript.

## Acknowledgements

Cermaq Norway AS, Hopen, Bodø is acknowledged for providing the fish at the freshwater stages. We thank Hilde Ribe and Katrine Klippenberg for their help in procuring the samples. Vigdis Edvardsen is acknowledged for her assistance while sequencing the libraries.

## Additional files

***Additional file 1:*** Additional methods

***Additional file 2:*** Fig. S1. Rarefaction curves based on the alpha diversity measure (PD Whole tree), for individual samples (a) and each stage (b). The curves indicate that a sequence number of 2400/sample is sufficient to capture most of the alpha diversity present in the samples as the curves become asymptotic at this depth.

***Additional file 2:*** Fig. S2. Plots showing the comparisons between the HL and 7wph. A stage-specific colour coding is used for figures a, b, c, f. (a) Alpha diversity indices (Shannon index, PD whole tree, Simpson’s evenness) of the bacterial communities and (b, c) UniFrac distances-based PCoA. Mean relative abundance of the 10 most abundant OTUs and their (d) phylum-level and (e) genus-level taxonomic ranks. Taxonomic classification-specific color coding is used in figures d, e, and the OTUs without any assignment are shown in grey. (f) Cladogram showing the significantly abundant taxonomic members in the different stages.

***Additional file 3:*** Fig. S3. Plots showing the comparisons between the stages 12 and 20wph. A stage-specific colour coding is used for figures a, b, c, f. (a) Alpha diversity indices (Shannon index, PD whole tree, Simpson’s evenness) of the bacterial communities and (b, c) UniFrac distances-based PCoA. Mean relative abundance of the 10 most abundant OTUs and their (d) phylum-level and (e) genus-level taxonomic ranks. Taxonomic classification-specific color coding is used in figures d, e, and the OTUs without any assignment are shown in grey. (f) Cladogram showing the significantly abundant taxonomic members in the different stages.

***Additional file 3:*** Fig. S4. Plots showing the comparison between stages 62wph - freshwater and 65wph - seawater. A stage-specific colour coding is used for figures a, b, c, f. (a) Alpha diversity indices (Shannon index, PD whole tree, Simpson’s evenness) of the bacterial communities and (b, c) UniFrac distances-based PCoA. Mean relative abundance of the 10 most abundant OTUs and their (d) phylum-level and (e) genus-level taxonomic ranks. Taxonomic classification-specific color coding is used in figures d, e, and the OTUs lacking taxonomy assignment are shown in grey. (f) Cladogram showing the significantly abundant taxonomic groups in each of the stages.

***Additional file 3:*** Fig. S5. Mean weighted Nearest Sequenced Taxon Index (NSTI) for the predicted metagenomes of the microbiota associated with the different stages. The NSTI scores were around 0.1 until the 20wph followed by a pronounced increase at 44wph and the values remained higher than 0.15 for the succeeding stages sampled, indicating increasing dissimilarity between the metagenome and available reference genomes.

***Additional file 4:*** Table S1: Sample metadata

***Additional file 5:*** Table S2: Read statistics concerning different samples

***Additional file 6:*** Table S3. ANOSIM comparisons and the corresponding p and R values for each of the comparisons

***Additional file 7:*** Table S4. The list of taxonomic features belonging to different groups with their corresponding p values and the LDA effect size

***Additional file 8:*** Table S5a. List of the 5 most abundance (>2.5%) KEGG modules that associated with different stages of development; Table S5b. List of KEGG modules that were significantly associated with different groups within each of the stages of development. Features passing the p-value filter 0.05 and the effect size filter 0.75 are listed

***Additional file 9:*** Table S6. ANOSIM comparisons and the corresponding p and R values for each of the comparisons

## References

1. Mach N, Berri M, Estellé J, Levenez F, Lemonnier G, Denis C et al. Early-life establishment of the swine gut microbiome and impact on host phenotypes. Environ Microbiol Rep. 2015;7(3): 554–69. doi:10.1111/1758-2229.12285.

2. Sommer F, Bäckhed F. The gut microbiota — masters of host development and physiology. Nat Rev Microbiol. 2013;11(4): 227–38. doi:10.1038/nrmicro2974.

3. Ye L, Amberg J, Chapman D, Gaikowski M, Liu W-T. Fish gut microbiota analysis differentiates physiology and behavior of invasive Asian carp and indigenous American fish. ISME J. 2014;8(3): 541–51. doi:10.1038/ismej.2013.181.

4. Avella MA, Place A, Du S-J, Williams E, Silvi S, Zohar Y et al. *Lactobacillus rhamnosus* accelerates zebrafish backbone calcification and gonadal differentiation through effects on the GnRH and IGF systems. PLoS ONE. 2012;7(9):e45572–e. doi:10.1371/journal.pone.0045572.

5. Arrieta M-C, Stiemsma LT, Amenyogbe N, Brown EM, Finlay B. The intestinal microbiome in early life: health and disease. Front Immunol. 2014;5:427-. doi:10.3389/fimmu.2014.00427.

6. Matamoros S, Gras-Leguen C, Le Vacon F, Potel G, de La Cochetiere M-F. Development of intestinal microbiota in infants and its impact on health. Trends Microbiol. 2013;21(4): 167–73. doi:10.1016/j.tim.2012.12.001.

7. Rodríguez JM, Murphy K, Stanton C, Ross RP, Kober OI, Juge N et al. The composition of the gut microbiota throughout life, with an emphasis on early life. Microb Ecol Health Dis. 2015;26:26050-.

8. Yatsunenko T, Rey FE, Manary MJ, Trehan I, Dominguez-Bello MG, Contreras M et al. Human gut microbiome viewed across age and geography. Nature. 2012;486(7402):222–7. doi:10.1038/nature11053.

9. Bakke I, Coward E, Andersen T, Vadstein O. Selection in the host structures the microbiota associated with developing cod larvae (*Gadus morhua*). Environ Microbiol. 2015;17(10): 3914–24. doi:10.1111/1462-2920.12888.

10. Forberg T, Sjulstad EB, Bakke I, Olsen Y, Hagiwara A, Sakakura Y et al. Correlation between microbiota and growth in mangrove killifish (*Kryptolebias marmoratus*) and Atlantic cod (*Gadus morhua*). Sci Rep. 2016;6:21192-. doi:10.1038/srep21192.

11. Stephens WZ, Burns AR, Stagaman K, Wong S, Rawls JF, Guillemin K et al. The composition of the zebrafish intestinal microbial community varies across development. ISME J. 2015;10(3): 644–54. doi:10.1038/ismej.2015.140.

12. Wong S, Stephens WZ, Burns AR, Stagaman K, David LA, Bohannan BJM et al. Ontogenetic differences in dietary fat influence microbiota assembly in the zebrafish gut. mBio. 2015;6(5):e00687–15. doi:10.1128/mBio.00687-15.

13. Llewellyn MS, McGinnity P, Dionne M, Letourneau J, Thonier F, Carvalho GR et al. The biogeography of the Atlantic salmon (Salmo salar) gut microbiome. ISME J. 2016;10(5): 1280–4. doi:10.1038/ismej.2015.189.

14. Yan Q, Li J, Yu Y, Wang J, He Z, Van Nostrand JD et al. Environmental filtering decreases with fish development for the assembly of gut microbiota. Environ Microbiol. 2016. doi:10.1111/1462-2920.13365.

15. Hansen GH, Olafsen JA. Bacterial colonization of cod (*Gadus morhua L*.) and halibut (*Hippoglossus hippoglossus*) eggs in marine aquaculture. Appl Environ Microbiol. 1989;55(6): 1435–46.

16. Llewellyn MS, Boutin S, Hoseinifar SH, Derome N. Teleost microbiomes: the state of the art in their characterization, manipulation and importance in aquaculture and fisheries. Front Microbiol. 2014;5:207-. doi:10.3389/fmicb.2014.00207.

17. Fujimoto M, Crossman JA, Scribner KT, Marsh TL. Microbial community assembly and succession on lake sturgeon egg surfaces as a function of simulated spawning stream flow rate. Microb Ecol. 2013;66(3): 500–11. doi:10.1007/s00248-013-0256-6.

18. Yoshimizu M, Kimura T, Sakai M. Microflora of the embryo and the fry of salmonids. Nippon Suisan Gakkaishi. 1980;46: 967–75. doi:10.2331/suisan.46.967.

19. Burns AR, Stephens WZ, Stagaman K, Wong S, Rawls JF, Guillemin K et al. Contribution of neutral processes to the assembly of gut microbial communities in the zebrafish over host development. ISME J. 2016;10(3): 655–64. doi:10.1038/ismej.2015.142.

20. Trust TJ. Sterility of salmonid roe and practicality of hatching gnotobiotic salmonid fish. Appl Microbiol. 1974;28(3): 340–1.

21. Zapata A, Diez B, Cejalvo T, Gutiérrez-de Frías C, Cortés A. Ontogeny of the immune system of fish. Fish Shellfish Immunol. 2006;20(2): 126–36. doi:10.1016/j.fsi.2004.09.005.

22. Liu Y, de Bruijn I, Jack ALH, Drynan K, van den Berg AH, Thoen E et al. Deciphering microbial landscapes of fish eggs to mitigate emerging diseases. ISME J. 2014;8(10): 2002–14. doi:10.1038/ismej.2014.44.

23. Rawls JF, Samuel BS, Gordon JI. Gnotobiotic zebrafish reveal evolutionarily conserved responses to the gut microbiota. Proc Natl Acad Sci USA. 2004;101(13): 4596–601. doi:10.1073/pnas.0400706101.

24. Gorodilov YN. Description of the early ontogeny of the Atlantic salmon, Salmo salar, with a novel system of interval (state) identification. Environ Biol Fishes. 1996;47(2): 109–27. doi:10.1007/BF00005034.

25. Bates JM, Mittge E, Kuhlman J, Baden KN, Cheesman SE, Guillemin K. Distinct signals from the microbiota promote different aspects of zebrafish gut differentiation. Dev Biol. 2006;297(2): 374–86. doi:10.1016/j.ydbio.2006.05.006.

26. Chung H, Pamp SJ, Hill JA, Surana NK, Edelman SM, Troy EB et al. Gut immune maturation depends on colonization with a host-specific microbiota. Cell. 2012;149(7): 1578–93. doi:10.1016/j.cell.2012.04.037.

27. Ingerslev HC, von Gersdorff Jørgensen L, Lenz Strube M, Larsen N, Dalsgaard I, Boye M et al. The development of the gut microbiota in rainbow trout (*Oncorhynchus mykiss*) is affected by first feeding and diet type. Aquaculture. 2014;424–425:24-34. doi:10.1016/j.aquaculture.2013.12.032.

28. Wells P, Pinder A. The respiratory development of Atlantic salmon. I. Morphometry of gills, yolk sac and body surface. J Exp Biol. 1996;199(Pt 12):2725–36.

29. Givens CE, Ransom B, Bano N, Hollibaugh JT. Comparison of the gut microbiomes of 12 bony fish and 3 shark species. Mar Ecol Prog Ser. 2015;518:209–23 doi:10.3354/meps11034.

30. Wong S, Waldrop T, Summerfelt S, Davidson J, Barrows F, Kenney PB et al. Aquacultured rainbow trout (*Oncorhynchus mykiss*) possess a large core intestinal microbiota that is resistant to variation in diet and rearing density. Appl Environ Microbiol. 2013;79(16): 4974–84. doi:10.1128/AEM.00924-13.

31. van Kessel MA, Dutilh BE, Neveling K, Kwint MP, Veltman JA, Flik G et al. Pyrosequencing of 16S rRNA gene amplicons to study the microbiota in the gastrointestinal tract of carp (*Cyprinus carpio* L.). AMB Express. 2011;1:41-. doi:10.1186/2191-0855-1-41.

32. Roeselers G, Mittge EK, Stephens WZ, Parichy DM, Cavanaugh CM, Guillemin K et al. Evidence for a core gut microbiota in the zebrafish. ISME J. 2011;5(10):1595–608. doi:10.1038/ismej.2011.38.

33. Holben WE, Williams P, Gilbert MA, Saarinen M, Särkilahti LK, Apajalahti JHA. Phylogenetic analysis of intestinal microflora indicates a novel *Mycoplasma* phylotype in farmed and wild salmon. Microb Ecol. 2002;44(2): 175–85. doi:10.1007/s00248-002-1011-6.

34. Lowrey L, Woodhams DC, Tacchi L, Salinas I. Topographical mapping of the rainbow trout (*Oncorhynchus mykiss*) microbiome reveals a diverse bacterial community with antifungal properties in the skin. Appl Environ Microbiol. 2015;81(19): 6915–25. doi:10.1128/AEM.01826-15.

35. Lokesh J, Kiron V. Transition from freshwater to seawater reshapes the skin-associated microbiota of Atlantic salmon. Sci Rep. 2016;6. doi:10.1038/srep19707.

36. Zarkasi KZ, Abell GCJ, Taylor RS, Neuman C, Hatje E, Tamplin ML et al. Pyrosequencing-based characterization of gastrointestinal bacteria of Atlantic salmon (*Salmo salar* L.) within a commercial mariculture system. J Appl Microbiol. 2014;117(1): 18–27. doi:10.1111/jam.12514.

37. Zarkasi KZ, Taylor RS, Abell GCJ, Tamplin ML, Glencross BD, Bowman JP. Atlantic salmon (*Salmo salar L*.) gastrointestinal microbial community dynamics in relation to digesta properties and diet. Microb Ecol. 2016;71(3): 589–603. doi:10.1007/s00248-015-0728-y.

38. Schmidt VT, Smith KF, Melvin DW, Amaral-Zettler LA. Community assembly of a euryhaline fish microbiome during salinity acclimation. Mol Ecol. 2015;24(10): 2537–50. doi:10.1111/mec.13177.

39. Sullam KE, Essinger SD, Lozupone CA, O’Connor MP, Rosen GL, Knight R et al. Environmental and ecological factors that shape the gut bacterial communities of fish: a meta-analysis. Mol Ecol. 2012;21(13): 3363–78. doi:10.1111/j.1365-294X.2012.05552.x.

40. Lozupone CA, Stombaugh JI, Gordon JI, Jansson JK, Knight R. Diversity, stability and resilience of the human gut microbiota. Nature. 2012;489(7415):220–30. doi:10.1038/nature11550.

41. Kozich JJ, Westcott SL, Baxter NT, Highlander SK, Schloss PD. Development of a dual-index sequencing strategy and curation pipeline for analyzing amplicon sequence data on the MiSeq Illumina sequencing platform. Appl Environ Microbiol. 2013;79(17): 5112–20. doi:10.1128/AEM.01043-13.

42. Edgar RC. UPARSE: highly accurate OTU sequences from microbial amplicon reads. Nat Methods. 2013;10(10): 996–8. doi:10.1038/nmeth.2604.

43. Edgar RC, Haas BJ, Clemente JC, Quince C, Knight R. UCHIME improves sensitivity and speed of chimera detection. Bioinformatics (Oxford, England). 2011;27(16): 2194–200. doi:10.1093/bioinformatics/btr381.

44. Caporaso JG, Kuczynski J, Stombaugh J, Bittinger K, Bushman FD, Costello EK et al. QIIME allows analysis of high-throughput community sequencing data. Nat Methods. 2010;7(5): 335–6. doi:10.1038/nmeth.f.303.

45. Segata N, Izard J, Waldron L, Gevers D, Miropolsky L, Garrett WS et al. Metagenomic biomarker discovery and explanation. Genome Biol. 2011;12(6):R60–R. doi:10.1186/gb-2011-12-6-r60.

46. McMurdie PJ, Holmes S. phyloseq: an R package for reproducible interactive analysis and graphics of microbiome census data. PloS one. 2013;8(4):e61217–e. doi:10.1371/journal.pone.0061217.

47. Asnicar F, Weingart G, Tickle TL, Huttenhower C, Segata N. Compact graphical representation of phylogenetic data and metadata with GraPhlAn. PeerJ. 2015;3:e1029–e. doi:10.7717/peerj.1029.

48. Langille MGI, Zaneveld J, Caporaso JG, McDonald D, Knights D, Reyes JA et al. Predictive functional profiling of microbial communities using 16S rRNA marker gene sequences. Nat Biotechnol. 2013;31(9): 814–21. doi:10.1038/nbt.2676.

49. Abubucker S, Segata N, Goll J, Schubert AM, Izard J, Cantarel BL et al. Metabolic reconstruction for metagenomic data and its application to the human microbiome. PLoS Comp Biol. 2012;8(6):e1002358–e. doi:10.1371/journal.pcbi.1002358.

50. Parks DH, Tyson GW, Hugenholtz P, Beiko RG. STAMP: statistical analysis of taxonomic and functional profiles. Bioinformatics (Oxford, England). 2014;30(21): 3123–4. doi:10.1093/bioinformatics/btu494.

